# A novel membrane stress-response that blocks chromosomal replication by targeting the DnaA initiator via the ClpP protease

**DOI:** 10.1101/2024.12.12.628197

**Authors:** Alabi Gbolahan, Tong Li, Rishit Saxena, Karen Wolcott, Aamna Sohail, Ishika Ahmad, Dhruba K. Chattoraj, Elliott Crooke, Rahul Saxena

## Abstract

In *Escherichia coli*, membrane-stress due to interrupted lipoprotein (Lpp) maturation impairs DNA replication and arrests cell growth. How the two disparate processes of Lpp maturation and DNA replication are connected remains unclear. We demonstrate that upon membrane-stress the Rcs stress-response pathway is activated and the replication initiator DnaA is lost, which explains the replication block. However, the Lon protease, a key regulator of the Rcs pathway, is not required for the DnaA loss. We further ruled out the involvement of (p)ppGpp, one of the major mediators of stress-response in bacteria. However, upon deletion of the ClpP protease gene, DnaA was stable, replication initiated, and there was no cell-growth arrest. In wildtype cells, overexpression of DnaA was lethal even without the membrane-stress apparently from hyper-initiation. The hyper-initiation was restrained in Δ*crp* cells, and overexpression of DnaA was able to overcome the growth-arrest. Δ*fis* cells, which were earlier found resistant to the membrane-stress, showed DnaA stability and normal replication upon induction of the membrane-stress. We conclude that DnaA loss suffices to explain the growth-arrest upon the membrane-stress. The stress-response pathway described here appears novel because of its independence from Lon and (p)ppGpp, which have been implicated in other stress-responses that block DNA replication.

**Significance:** The seminal observation that DNA replication-stress can block cell division in *E. coli* (SOS response), introduced the concept of checkpoint control in the cell cycle. Here, we describe a novel checkpoint control that functions in the opposite direction: membrane-stress causing replication block. We show how two apparently unrelated outcomes of reducing a membrane phospholipid, accumulation of a precursor lipoprotein (pLpp) and block of replication initiation, could be linked. The pLpp accumulation stresses the membrane that causes a response culminating in activating the ClpP protease that blocks replication by targeting the initiator DnaA. DnaA being vital and highly conserved, the detail understanding of the response pathway is likely to open new avenues to treat bacterial infection.

## Introduction

In Gram-negative bacteria such as *Escherichia coli*, the cell envelope consists of two concentric lipid bilayers, the outer membrane (OM) and the inner membrane (IM) with a periplasmic space in between wherein lies the peptidoglycan layer, the cell wall (Fig. 1). The OM has lipid-attached proteins (Lipoproteins) that are an evolutionarily conserved family of acylated proteins. These serve several functions, including biogenesis of membranes and sensing of stress from external or internal stimuli (1, 2). In *E coli*, the most abundant lipoprotein, Lpp (∼10^6^ molecules per cell), provides structural support to the OM by crosslinking it to the peptidoglycan layer (3, 4). The synthesis of nascent Lpp (product of the *lpp* gene) occurs in the cytosol as prelipoprotein (pLpp), which locates to the IM (1–4). In the IM, pLpp is modified at its Cys21 residue by the addition of diacylglycerol (DG) from phosphatidylglycerol (PG), the major anionic phospholipid of the IM (5–8). This lipidation of pLpp is catalyzed by diacylglycerol transferase, Lgt (9). The resultant DGLpp is cleaved by the IM-bound type II aspartyl endopeptidase, LspA (10). The cleaved product is processed further to produce the mature Lpp that translocates to the OM (1-8 and Fig. 1). Although Lpp provides structural support to the OM by crosslinking it to the peptidoglycan layer, Δ*lpp* mutants are viable, indicating that Lpp is not essential (11). Nevertheless, interrupting processing of pLpp at various steps of its maturation results in the accumulation of intermediates in the IM, and *E. coli* becomes non-viable (9-14 and Fig. 1).

**Fig. 1.**
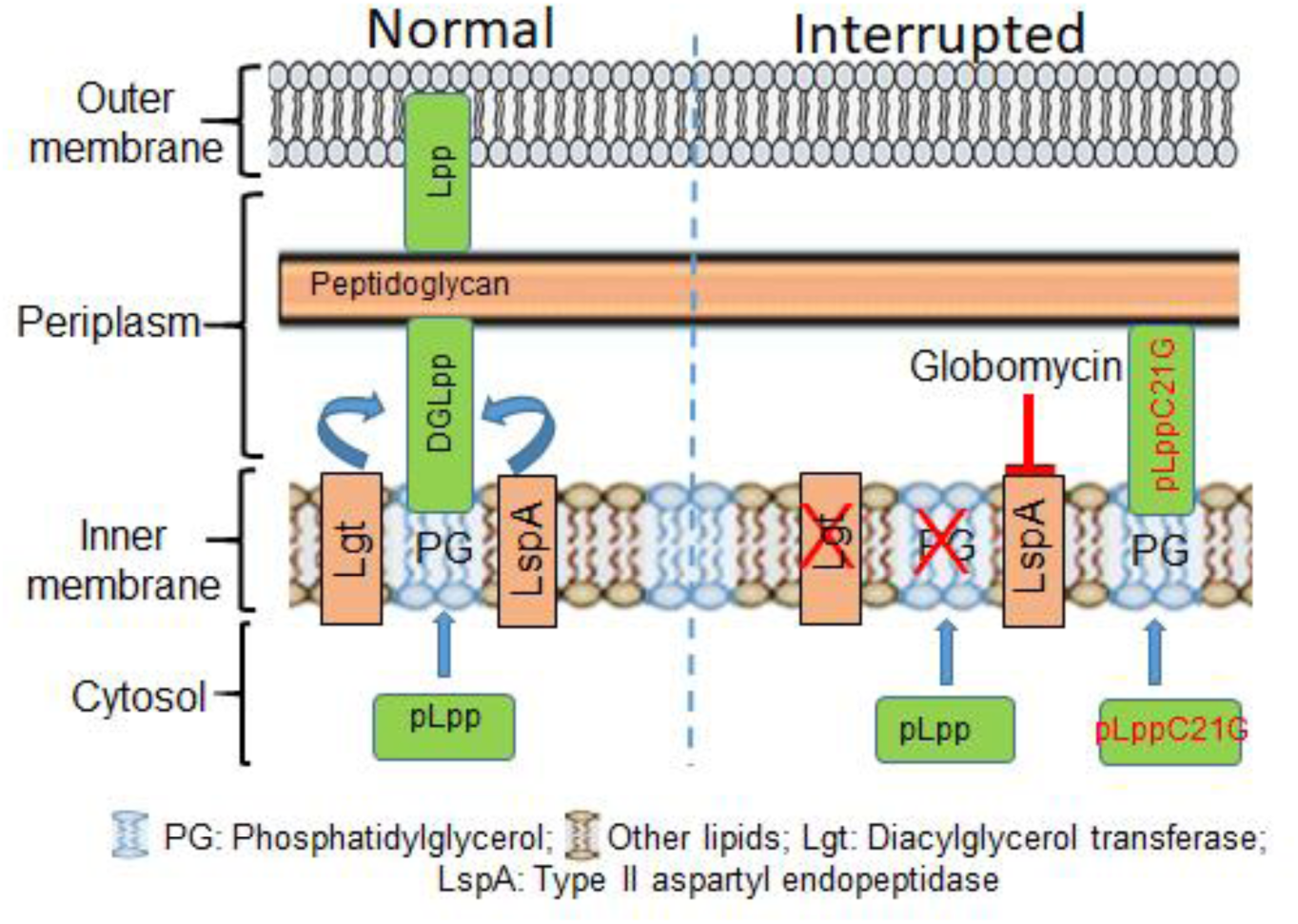
Schematic representation of Lpp maturation pathway and how it is interrupted at different steps. Lpp is synthesized as pLpp, interacts with PG and gets modified by Lgt to DGLpp. DGLpp is processed by LspA and others, and mature Lpp is incorporated in the outer membrane. The maturation is interrupted (indicated in Red) either by mutating Lgt, depleting PG, inactivating LspA by Globomycin or expressing a processing defective *lpp*(C21G) mutant. The interrupted products accumulate in the inner membrane, causing membrane-stress.

An early indication of unprocessed pLpp accumulation causing lethality in *E. coli* came from the repression of the PG synthase A (*pgsA*) gene expression, which would prevent the DG modification step of pLpp (12, 13). These results were obtained by deleting the native *pgsA* gene and placing a copy of the gene ectopically under the IPTG-inducible P*_lac_* promoter. Removal of IPTG from a fully induced culture caused a 10-fold reduction in PG level and cell growth-arrest (12, 13). Cells under these conditions are referred to as “*pgsA*-null”. Adding back IPTG to the growth medium restored cell growth, indicating that the growth-arrest was due to repression of *pgsA* expression (12, 13). Deletion of the *lpp* gene of *pgsA*-null cells allowed their growth also (15, 16). These results indicate that the growth-arrest of *E. coli* containing reduced PG levels could be associated with defective Lpp biogenesis. The Lpp biogenesis in wildtype *E. coli* (*pgsA^+^*) could also be interrupted: by deleting the *lgt* gene (9) or overexpressing mutant pLpp(C21G), either of which would interfere with the DG modification step (11, 14), or by blocking LspA enzyme activity, which cleaves the IM-bound DGLpp, using a lipopeptide inhibitor, globomycin (17, 18) [Summarized in Fig. 1]. In sum, membrane-stress from interrupted lipoprotein biogenesis in any of the steps of the pLpp maturation pathway can cause growth-arrest but the mechanistic basis of the arrest remains unclear.

The growth-arrested *pgsA-*null cells also have decreased frequency of replication initiation (19, 20). PG depletion thus causes blockage of two apparently unrelated processes: Lpp maturation and chromosomal DNA replication (14). Here, we have investigated how these two processes could be linked.

Initiation of chromosomal replication in *E. coli* occurs when multiple DnaA molecules oligomerize at a unique 245 base pair origin of replication (*oriC*) (21–23). Wrapping of the origin DNA around an oligomer of DnaA protein together with HU protein and near physiological concentrations of ATP allows opening of an *oriC* stretch, called the DNA unwinding element (DUE) (21–23). ATP-DnaA but not ADP-DnaA, produces replication-proficient *oriC*-DnaA complexes required for the DUE opening (23–27). Once DUE is unwound, DnaB helicase and, subsequently, the rest of the replisome components can be loaded onto the origin so that DNA synthesis can commence (28).

Biochemical studies have shown that membrane lipids containing anionic head groups, like PG and CL (cardiolipin), can rejuvenate inactive initiator DnaA-ADP to active initiator DnaA-ATP (30, 31). Imaging studies have indicated that DnaA is associated peripherally to the membrane (32–34). In fact, DnaA has a stretch of amino acid residues in its ATPase (AAA^+^) domain that can bind membranes (35, 36). That the membrane state might be involved in DnaA function is suggested by the finding that the overexpression of DnaA(L366K), where the changed residue is in the membrane-binding stretch of the AAA+ domain, in combination with DnaA(WT), but not overexpression of DnaA(WT) alone, suppresses growth-arrest due to defective Lpp maturation [as in *pgsA-*null cells or wildtype (*pgsA*^+^) cells overexpressing pLpp(C21G)] (14,35–37). Similarly, we have recently shown that overexpression of DnaA with small deletions in its linker domain, a domain that also has membrane-binding capacity, can overcome the growth-arrest (38). These results suggest that the growth-arrest from defective Lpp biogenesis is mediated through DnaA, possibly by blocking replication initiation.

Transient faults in several cellular processes often induce a response (stress-response), which buys time for cells to adapt to stressful conditions. A well-studied stress-response is mediated via the Rcs (Regulator of capsule synthesis) pathway, which is activated primarily due to defects in lipopolysaccharide (LPS) synthesis (39–41). The activation of the pathway is known in *pgsA*-null cells (42). In the Rcs system, the phosphate transfers from RcsC (the sensor kinase) to RcsD (the phosphate transmitter), both components of the IM, and finally to RcsB, the cytoplasmic DNA-binding protein (response regulator) (39–41). RcsB binds to RcsA and the RcsB-RcsA dimer induces transcription of *cps* genes that control colonic acid production (the main component of capsular polysaccharides attached to the OM), making *E. coli* cells mucoid (39–41). The elevated capsular polysaccharide level is considered the hallmark of Rcs stress-response. The three positive regulators, RcsA, RcsB and RcsF, and two negative regulators, RcsC and Lon (39, 40) control the Rcs stress-response. The Lon protease degrades RcsA to downregulate capsular polysaccharide synthesis under normal growth (40). Δ*lon* cells are mucoid due to increased *cps* gene expression apparently from unchecked RcsA level in the absence of Lon (41).

In *pgsA*-null cells, induction of the Cpx system is also known (42). The system comprises an OM stress-receptor protein, NlpE, a sensor kinase CpxA, and the response regulator CpxR that binds to RNAP (43, 44). The CpxAR system monitors the trafficking of OM-targeted lipoproteins and responds to their mislocalization in IM *via* activation of the DegP protease (43, 44). The extracytoplasmic stress could also be relieved by activation of ClpP (caseinolytic protease P). ClpP functions with ATPases, ClpA or ClpX by forming ClpAP or ClpXP heterodimers (44, 45), which can degrade damaged, misfolded, and regulatory proteins and thus help maintain protein quality control (44).

In several other stresses, synthesis of nucleotide-based second messengers is induced, that is required to cause the protease-dependent response. Such messengers include guanosine tetraphosphate (ppGpp) and guanosine pentaphosphate (pppGpp), collectively known as (p)ppGpp, the hallmark of stringent response in bacteria (46). (p)ppGpp synthesis is induced in bacteria under various stressful conditions, such as nutrient limitation, oxygen free-radical production, fatty acid limitation, and antibiotic treatment (47–49). ppGpp inhibits the activity of PPX (exopolyphosphatase), which hydrolyzes polyphosphate (PolyP) (46). Accumulation of PolyP in bacteria may activate the Lon protease (50), which can degrade the DnaA protein (51, 52). (p)ppGpp is known mainly for reprogramming transcription and blocking translation (53). (p)ppGpp either binds directly to RNA polymerase (RNAP) or regulates the activity of the housekeeping Sigma factor, σ^70^ to alter RNAP interaction with many promoters (54). (p)ppGpp can also inhibit initiation of DNA replication (51, 52). (p)ppGpp activates the Lon protease that selectively degrades ADP-DnaA (52), which occupy *dnaA* promoters (25) or DnaA reactivating sites (RIDAs) on chromosomal DNA (55) to promote *de novo* synthesis of DnaA to produce replication-efficient ATP form or change transcription and thereby decreasing DNA superhelicity near *oriC* and inhibiting replication initiation (56).

Besides (p)ppGpp, bacteria can synthesize other nucleotide-based secondary messengers, including the cyclic (3′,5′)-adenosine phosphate, cAMP (57). Adenylyl cyclase (product of the *cyaA* gene) catalyzes the conversion of ATP to cAMP, which acts as an allosteric effector of the catabolite repressor protein, CRP (product of the *crp* gene), expressed during glucose starvation (58, 59). Like (p)ppGpp, cAMP-CRP complex directly contacts RNAP or regulates σ^70^ binding to RNAP to alter transcription of genes from various promoters, eventually controlling many cellular activities, including stress-response pathways (60, 61).

In this study, the growth-arrest of *E. coli* under membrane-stress due to interrupted Lpp-maturation appears to be due to ClpP-mediated proteolysis of DnaA and thereby blocking of new rounds of replication initiation. Overexpression of DnaA, although lethal in wildtype cells due to overinitiation was restrained in Δ*crp* cells. Under this condition, Δ*crp* cells could overcome the growth-arrest upon the membrane-stress. The growth-arrest was also overcome earlier without DnaA over-expression in Δ*fis* cells. In these cells, DnaA was found to be stable upon induction of the stress. These results demonstrate that DnaA loss is responsible for cell growth-arrest upon induction of the membrane stress. The delineation of the pathway from induction of the stress to the loss of DnaA promises to advance our understanding of replication regulation when cells are under stress and how the knowledge can be harnessed to arrest bacterial growth in infection.

## Results

### DnaA is lost in cells growth-arrested due to interrupted lipoprotein biogenesis

In *E. coli*, the requirement of PG is evident both for processing pLpp (8–11) and for initiation of chromosomal DNA replication (19, 20). We showed that as opposed to *lpp*(WT) expression of a mutant gene *lpp*(C21G), whose product cannot be processed and that accumulates in the IM, causes growth-arrest of wildtype *E. coli* cells (14). The growth-arrest could be overcome by overexpression of not the WT DnaA but a mutant DnaA(L366K), where the change is in one of the membrane-interacting interfaces of the initiator (14, 37, 38). These studies implied that the membrane stress-response causing the growth-arrest could be mediated through DnaA, but in what way remained an open question.

To address this, we used plasmids to overproduce Lpp(WT) and pLpp(C21G) proteins from the inducible P*_lac_* promoter by including the inducer IPTG in the growth medium (14). In addition to Lpp(WT), we used another negative control, the pLpp(C21G/ΔK) protein. Normally, pLpp covalently crosslinks with the peptidoglycan layer using the carboxy-terminal lysine, deletion of which keeps the mutant protein (pLpp(C21G/ΔK) cytosolic and thereby not causing membrane-stress and growth-arrest (62). Upon addition of 50 µM IPTG, cells expressing pLpp(C21G) protein grew up to OD_600_ of 0.10, whereas the cells expressing the negative controls Lpp(WT) and pLpp(C21G/ΔK) reached a higher OD_600_ of 0.3-0.5 in the same time interval (Fig. 2 A, *left panel*). The generation time also increased ∼1.4-fold when cells were expressing pLpp(C21G) as opposed to Lpp(WT) or pLpp(C21GΔK) (Supplementary Table S2). Immunoblotting data using polyclonal α-DnaA-antiserum suggest that DnaA protein is lost in cells expressing pLpp(C21G) and not in cells expressing Lpp(WT) or pLpp(C21G/ΔK) (Fig. 2 A, *right panel*). When pLpp maturation was blocked by using globomycin, the generation time also increased compared that of cells with no drugs (Fig. 2 B, *left panel* and Supplementary Table S2). We also confirmed that the globomycin-induced toxicity requires Lpp since *lpp*^+^ cells showed increased toxicity against globomycin as compared to *Δlpp* cells, as was suggested in a previous study (63, and Supplementary Fig. S1A). The immunoblotting data showed that as in the wildtype (*pgsA^+^lpp^+^*) cells expressing *lpp*(C21G), DnaA was lost when the wildtype cells were treated with globomycin (Fig. 2 B, *right panel*). The growth-arrest thus can be attributed to the DnaA loss. In the remainder of this study, membrane-stress will be studied mainly in wildtype (*pgsA^+^lpp^+^*) cells either by expressing plasmid-borne *lpp*(C21G) gene or by treatment with globomycin.

**Fig. 2.**
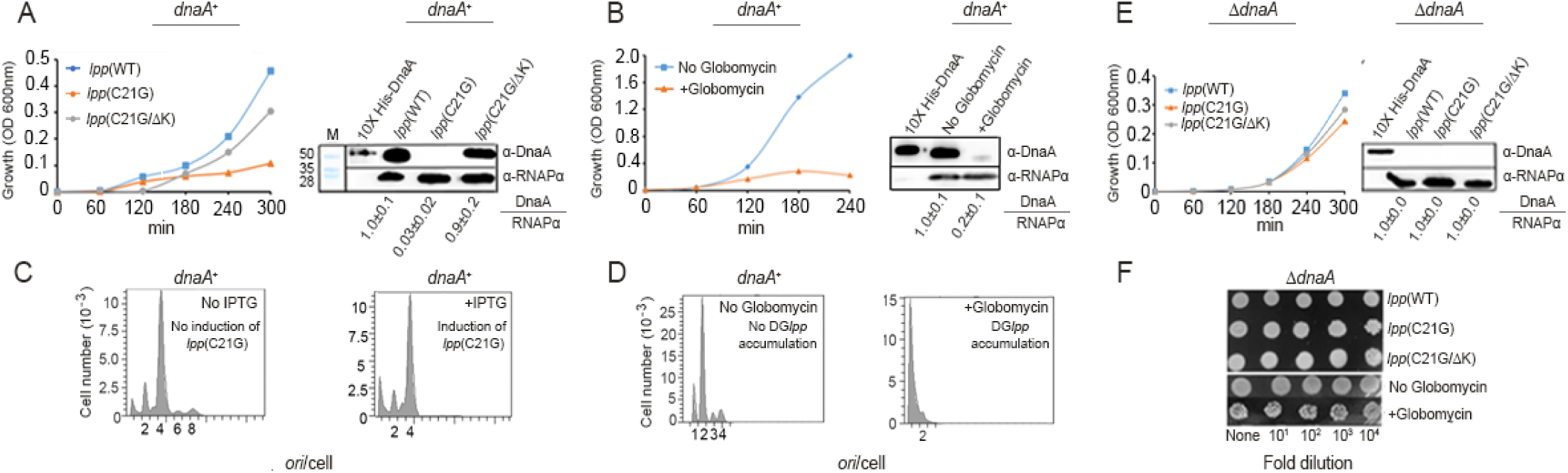
Membrane-stress upon interrupting Lpp maturation results in loss of DnaA initiator and block of replication initiation. (*A*) Growth of wildtype (*dnaA*^+^) cells expressing *lpp*(WT), *lpp*(C21G), and *lpp*(C21GΔK) genes (*left panel*). Immunoblotting (*right panel*) of DnaA (52 kDa) and α-subunit of RNA polymerase (28 kDa used as loading control) from cells used in the left panel. M indicates protein size marker lane. Purified 10X His-tagged DnaA was used as DnaA size marker. Numbers indicate the relative ratios of DnaA to RNAPα proteins. The DnaA/RNAPα ratios are normalized w.r.t. the *lpp*(WT) value of 1.0. (*B*) Growth of wildtype cells in the absence and presence of globomycin (*left panel*). Immunoblotting of DnaA and RNAPα from such cells (*right panel*). Other details are same as in (*A*). (*C-D*): Replication run-off profiles after rifampicin addition analyzed by flow cytometry. Numbers in abscissa indicate chromosomal origins per cell at the time of rifampicin addition. (*E*) Growth of Δ*dnaA* cells expressing *lpp*(WT), *lpp*(C21G), and *lpp*(C21GΔK) genes (*left panel*) and immunoblotting of DnaA and RNAPα from such cells (*right panel*). (*F*) Spotting assay to test viability of Δ*dnaA* cells expressing *lpp*(WT), *lpp*(C21G) or *lpp*(C21GΔK) genes or when the cells were exposed or not to globomycin.

The loss of DnaA under the membrane-stress is expected to affect replication initiation frequency at the chromosomal *oriC*. This was tested by flow cytometry. When *lpp*(C21G) was expressed in wildtype cells or when the cells were treated with globomycin new rounds of replication were blocked in both the cases (Fig. 2 C and D, *left vs right panel*). Note that most of the cell population carries two to four origins per cell even after induction of *lpp*(C21G) expression (Fig. 2C *right panel*). This we believe could be because it takes time for the unprocessed lipoprotein intermediates to accumulate to a threshold level before they become inhibitory. However, blockage of new rounds of replication was still conspicuous by the absence of 6 and 8 *ori-*carrying cells (Fig. 2 C, *right panel*). Globomycin was more effective in blocking replication, apparently because the drug action was more immediate than that could be achieved in an inducible system, where the inhibitor needed to be synthesized and accumulated (Fig. 2D, *left vs right panel*). These results indicate that the interrupted trafficking of a nonessential lipoprotein, Lpp, causes DnaA loss, blocking the vital event of replication initiation.

The *oriC* region and the *dnaA* gene although normally are essential for initiation of replication, their requirement can be bypassed by other means (64–66). For example, *E. coli* Δ*dnaA* cells can be made viable by integrating into the chromosome a miniR1 plasmid (pKN500), where the replication initiates from the plasmid origin independently of DnaA (66). We asked whether in this plasmid-integrated Δ*dnaA* cells, the growth-arrest in response to membrane-stress could be bypassed, if the growth arrest were due to blockage of DnaA-mediated replication initiation. We transformed the Δ*dnaA* cells with plasmids expressing Lpp or its mutants and confirmed by immunoblotting that no band corresponding to DnaA protein is present in any of the transformants of Δ*dnaA* cells (Fig. 2E, *right panel*). As expected, overexpression of *lpp*(C21G) did not affect the growth of Δ*dnaA* cells, unlike the situation in wildtype cells (Fig. 2E, *left panel* & 2F, *top panel*). Similarly, Δ*dnaA* cells were resistant to globomycin, as opposed to wildtype cells (Fig. 2F, bottom *panel*). In addition, in Δ*dnaA* cells, no significant differences in growth rate were found when the WT or mutant Lpp proteins were expressed (Supplementary Table S2). We conclude that the interrupted trafficking of a nonessential lipoprotein, Lpp, perturbs specifically the DnaA-dependent replication initiation.

### Membrane-stress activates the Rcs stress-response

Stresses to cells due to internal dysfunction or external insults can induce responses (stress-responses), which buys time to repair damages from the stress and help cells return to normalcy. One of the best-studied membrane stress-response is the Rcs (Regulator of capsule synthesis) pathway. This pathway involves transcriptional activation of a complex network of capsular polysaccharide (*cps*) genes controlling the production of colonic acid, which makes cells mucoid (39, 40). The protease Lon serves as the negative regulator of colonic acid production. Lon degrades RcsA, one of the subunits of the transcriptional activator RcsA-RcsB heterodimer, and thus inhibits expression of the *cps* genes and prevents cells from becoming mucoid. Δ*lon* cells are, therefore, mucoid (41).

When spread on agar plates containing IPTG (50 µM), we found that cells under membrane-stress, i.e., those expressing the plasmid-borne *lpp*(C21G) gene exhibit the mucoidy phenotype (Fig. 3A, *middle panel*). Mucoidy was not seen in cells expressing the negative controls, *lpp*(WT) and the *lpp*(C21G/ΔK) genes (Fig. 3A, *left* and *right panels*, respectively), suggesting that the mucoidy is specific to cells accumulating pLpp(C21G) in the membrane. The activation of the Rcs stress-response is known in *pgsA*-null cells, where pLpp maturation is expected to be blocked causing it to accumulate in the membrane and stressing it (67). To test whether the Rcs pathway is on in mucoid cells with accumulated pLpp(C21G), we isolated total RNA from these cells (collected from plates) and performed qRT-PCR analysis. We found that mucoid cells expressing *lpp*(C21G) have increased levels of mRNA belonging to *rcsA*, *cpsB,* and *cpsG* genes (Fig. 3B). These results confirm turning on of the Rcs pathway when cells are under stress due to *lpp*(C21G) expression.

**Fig. 3.**
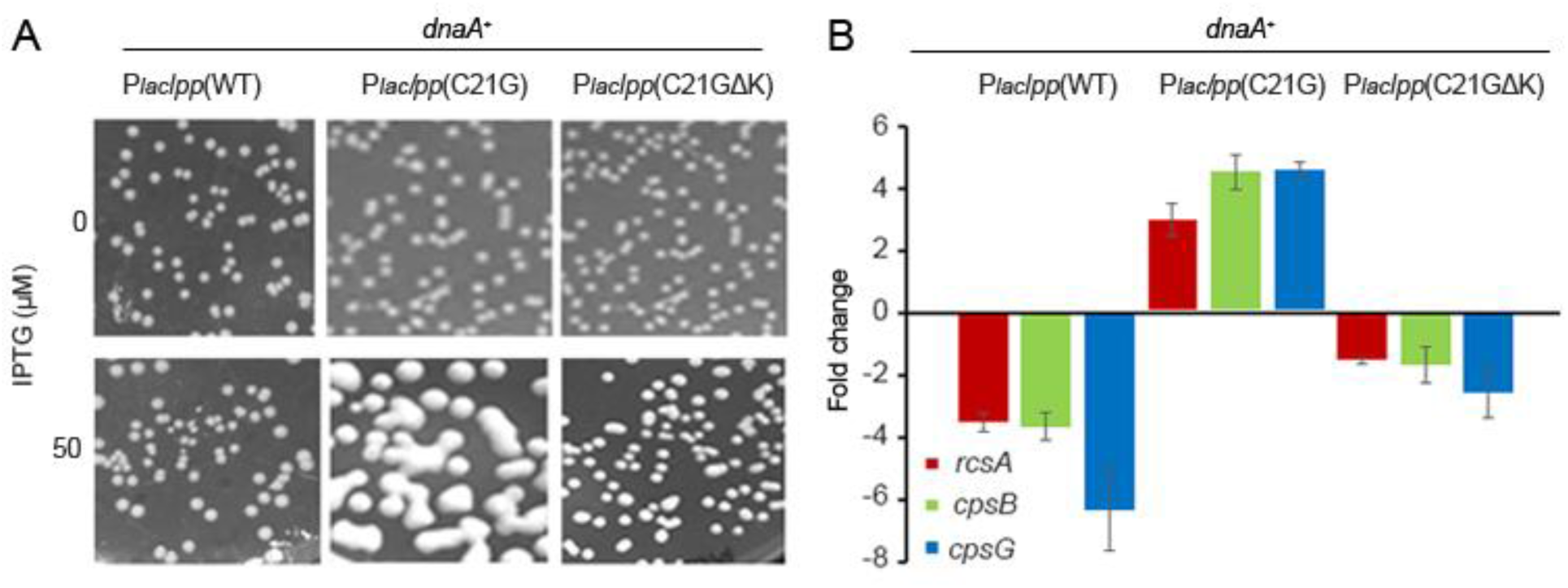
Membrane-stress upon interrupting Lpp maturation induces Rcs stress-response. Plasmids carrying *lpp*(WT), *lpp*(C21G) or *lpp*(C21GΔK) genes placed under the P*_lac_* promoter were used to transform wildtype (*dnaA^+^*) cells. (*A*) Growth phenotype of cells expressing WT and mutant Lpp genes in the presence of indicated amounts of IPTG. (*B*) qRT-PCR analysis of *rcsA*, *cpsB* and *cpsG* mRNA in cells expressing *lpp*(WT), *lpp*(C21G) or *lpp*(C21GΔK) genes. Data were normalized to the ΔCq value of a reference gene *rrsA*.

### Lon protease is not required for DnaA loss when cells are under membrane-stress

Our results so far indicated that the membrane-stress upon interrupted lipoprotein biogenesis activates the Rcs stress-response pathway and causes DnaA loss. However, the cause of the loss remains unknown. *E. coli* under stressful conditions, such as nutrient depletion, antibiotic treatment, and defective oxidative phosphorylation often accumulates (p)ppGpp molecules, a hallmark of the stringent response (46–51). (p)ppGpp activates the Lon protease in *C. crescentus* (68, 69) and *E. coli* (51, 52) that can degrade DnaA, blocking new rounds of replication. These results prompted us to investigate if blocking stringent-response or deleting the *lon* gene could allow cells to tolerate membrane-stress caused by *lpp*(C21G) expression.

To test this, we used *E. coli* Δ*relA* and Δ*lon* cells. In Δ*relA* cells, the stringent response is not induced because RelA is required for (p)ppGpp synthesis (70, 71). We transformed Δ*relA* and Δ*lon* cells with a plasmid containing the P*_lac_lpp*(C21G) gene. However, similar to wildtype cells, the growth for both Δ*relA* (Fig 4A, *left panel*) and Δ*lon* (Fig 4A, *right panel*) cells was inhibited when the inducer (50 µM IPTG) was included in the medium. Immunoblotting data showed that both Δ*relA* (Fig 4B, *top panel*) and Δ*lon* (Fig 4B, *bottom panel*) cells expressing *lpp*(C21G) have lost DnaA. The flow cytometry profile of such cells indicated a block of chromosomal replication initiation, as would be expected if DnaA were lost (Fig. 4C *left vs. right panels*). These results indicate that (p)ppGpp and Lon activities are not essential for the DnaA loss and blocking of replication due to interrupted lipoprotein maturation.

**Fig. 4.**
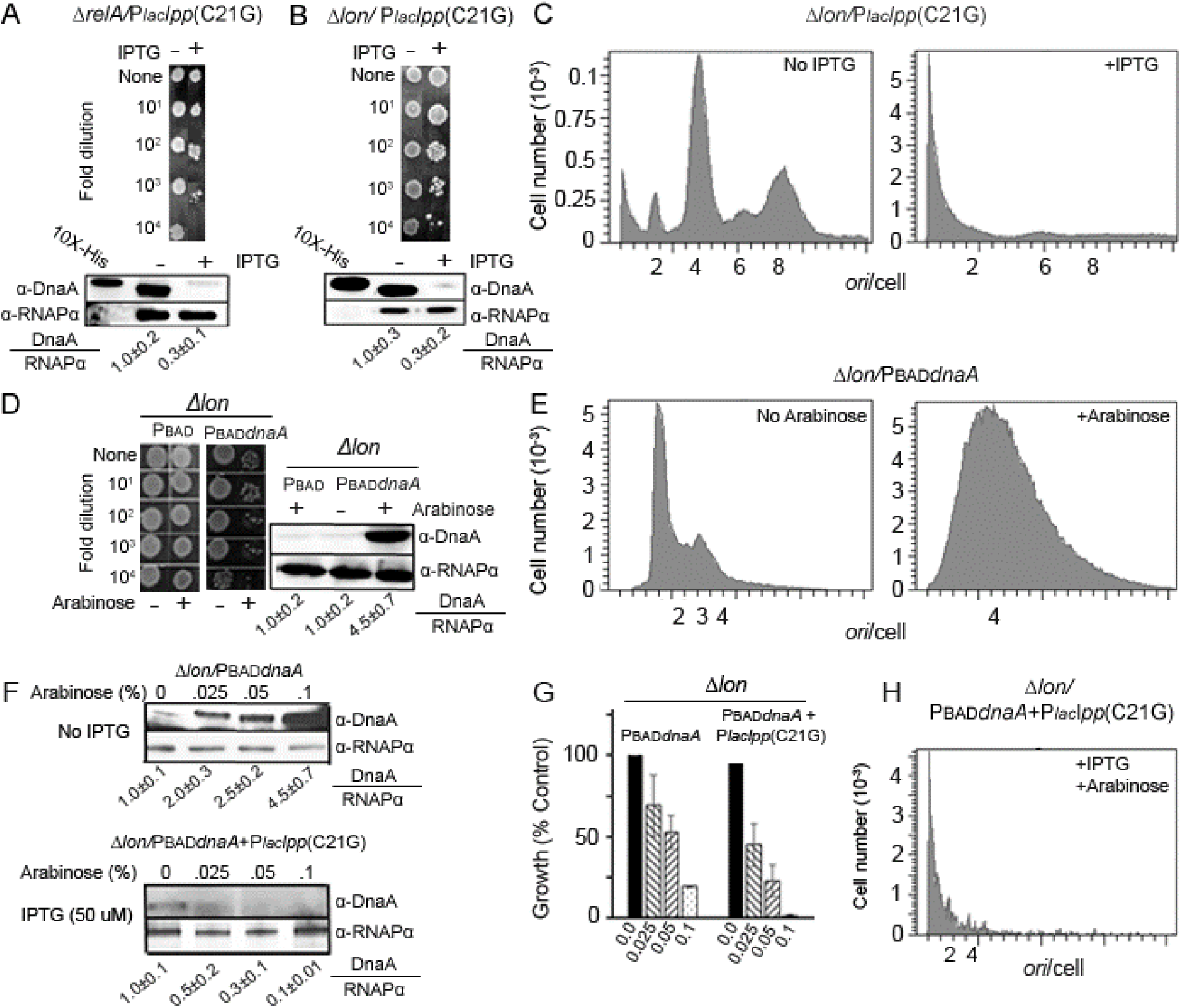
Depletion of DnaA upon membrane-stress is independent of (p)ppGpp alarmone and Lon protease, and is not overcome by overexpressing DnaA. *(A-B) E. coli* Δ*relA* (the gene required for (p)ppGpp synthesis) or Δ*lon* cells, transformed with plasmid carrying the stressor gene P*_lac_lpp*(C21G) were grown to OD_600_ of 0.10 and the cultures serially diluted. 2.5 µL from each dilution was spotted on M9-Agar plates without and with 50 µM IPTG to express the stressor gene (*top panels*). Immunoblotting to measure DnaA content of Δ*relA* and Δ*lon* cells expressing the stressor (*lower panels*). Other details are same as in Fig. 2A. (*C*) Replication run-off profiles of Δ*lon* cells carrying the stressor plasmid in the absence (*left panel*) and presence (*right panel*) of IPTG. *(D)* Properties of Δ*lon* cells carrying plasmids containing P_BAD_ or P_BAD_*dnaA*. Their growth in the absence (*left panel*) or presence (*middle panel*) of arabinose by spot assay and DnaA content by immunoblotting (*right panel*). (*E*) Origin content of the same cells as in (*C*) was measured by flow cytometry. (*F*) DnaA is depleted upon the membrane-stress even when over-expressed. Δ*lon* cells carrying the P_BAD_*dnaA* plasmid were grown to OD_600_ of 0.10 and plated on M9-Agar plates containing appropriate antibiotics and arabinose at various concentrations (0, 0.025, 0.05, and 0.1%). Single colonies from such plates were inoculated in liquid media with identical supplements and DnaA content of the cells were determined by immunoblotting (*top panel*). The Δ*lon/*P_BAD_*dnaA* cells were further transformed with the plasmid with the stressor gene [P*_lac_lpp*(C21G)] and were selected with appropriate antibiotics but without adding any inducer. The transformants were used to inoculate liquid media as before but in the absence or presence of 50µM IPTG before the cultures were probed for DnaA (*bottom panel*). Other details are same as in (Fig. 2A). (*G*) Cell viability of Δ*lon* cells expressing DnaA or co-expressing *dnaA* and *lpp*(C21G) genes. (*H*) Replication run-off profile of Δ*lon* cells co-expressing *dnaA* and *lpp*(C21G) genes, as in (*G*).

It has been recently reported that inhibition of new rounds of DNA replication due to stringent response is not solely due to the lowering of DnaA content, as overproduction of DnaA from plasmids does not remove the replication block (72). These results prompted us to examine whether in Δ*lon* cells, supplying extra DnaA from a plasmid source could compensate for the DnaA loss due to the membrane-stress. For this, Δ*lon* cells were transformed with plasmids carrying P_BAD_ and P_BAD_*dnaA*. As expected, in the absence of arabinose growth of bacteria containing P_BAD_ and P_BAD_*dnaA* plasmids but not the *lpp*(C21G)-carrying stressor plasmid, remained unaffected (Fig. 4D, *left panel*). However, when arabinose was present, growth of Δ*lon* cells was inhibited (Fig. 4D, *middle panel*). Immunoblotting confirmed the overproduction of the WT DnaA protein when the inducer was present (Fig. 4D, *right panel*). In flow-cytometric analysis, the replication initiation appeared excessive in Δ*lon* cells overproducing DnaA(WT) *versus* when the same cell carried the empty P_BAD_ plasmid (Fig. 4E, *left vs. right panels*). These results raised the possibility that the lethality due to DnaA(WT) overproduction in wildtype (14, and Supplementary Fig. S1B) and Δ*lon* cells (Fig. 4D) could be associated with hyper replication initiation.

An earlier report also suggested that overproduction of DnaA could be toxic to cells, although a limited increase in the amount of DnaA may be tolerated (72). This led us to test whether a more limited increase in DnaA, which by itself is not lethal, can bypass growth-arrest due to the membrane-stress. For this, Δ*lon* cells carrying P_BAD_*dnaA* plasmid were grown without arabinose up to OD_600_ of 0.1 and the cultures were spread on agar plates containing different amounts of arabinose. The cells collected from the plates were tested for the presence of DnaA by immunoblotting (Fig. 4F). The data confirmed overproduction of DnaA in the presence of arabinose (Fig. 4F, *top panel*). Increasing the DnaA expression in Δ*lon* cells did reduce growth (about four-fold at the highest inducer concentration of 0.1% (Fig. 4G, *left panel*). To test whether this level of overproduction could overcome the membrane-stress, the Δ*lon* cells carrying the P_BAD_*dnaA* plasmid were further transformed with the *lpp*(C21G)-carrying plasmid. Coexpression of *lpp*(C21G) and *dnaA* further reduced cell growth (Fig. 4G, *right panel*). Immunoblotting results indicated that in the presence of pLpp(C21G) even when *dnaA* expression was induced, DnaA was proteolysed (Fig. 4F, *bottom panel*). The flow cytometry results also indicated a severe loss of DNA upon induction of *lpp*(C21G) under conditions of *dnaA* overexpression (in the presence of 50 µM IPTG and 0.1 % arabinose) (Fig. 4H). Together, these results indicate that in the absence of membrane-stress overproduction of DnaA itself could be lethal due to hyper replication initiation, and upon the membrane-stress the lethality could be from DnaA proteolysis. In the latter case, DnaA was proteolysed even when overexpressed. Thus, attempts to overcome the growth-arrest in wildtype and Δ*lon* cells by overproducing DnaA proved futile, although for different reasons in the two cells.

### ClpP-deleted cells upon induction of the membrane stress do not lose DnaA and are not growth-arrested

Our results so far indicate that upon induction of the membrane-stress, DnaA is lost without requiring the Lon protease (Figs. 2 and 4). Bacteria can respond to the extracytoplasmic stress via activating in addition to Rcs, another two-component system, Cpx (44). The Cpx system controls the activation of the periplasmic DegP protease involved in the clearance of aggregated proteins from the periplasmic space and IM (73). Moreover, the cytoplasmic ClpP protease could also mitigate the stress upon accumulation of mislocalized proteins (74). To test the role of these proteases, we transformed *E. coli* Δ*degP* and Δ*clpP* cells with plasmid carrying the stressor P*_lac_lpp*(C21G) gene. Cell viability was tested with and without induction of the stressor gene. The Δ*degP* cells expressing the stressor did not grow (Supplementary Fig. S2). Immunoblotting showed loss of DnaA in such cells. The stress-response thus can happen without the DegP protease.

In contrast, we found that stress-induction is no longer lethal to Δ*clpP* cells (Fig. 5A, *left panel*). The result is consistent with the immunoblotting data, which showed the presence of DnaA upon stress-induction in Δ*clpP* cells (Fig. 5A, *right panel*). The flow cytometric analysis confirmed normal DNA replication in Δ*clpP* cells without and with stress-induction (Fig. 5B). Results were similar when the stress was induced using globomycin rather than expressing *lpp*(C21G) (Fig. 5C). As would be expected from DnaA stability, the Δ*clpP* cells were still susceptible to lethality from *dnaA* overexpression (Fig. 5D) and consequent hyper-initiation (Fig. 5E). In sum, it appears that induction of pLpp accumulation-mediated membrane-stress blocks cell growth by proteolysing DnaA that requires ClpP.

**Fig. 5.**
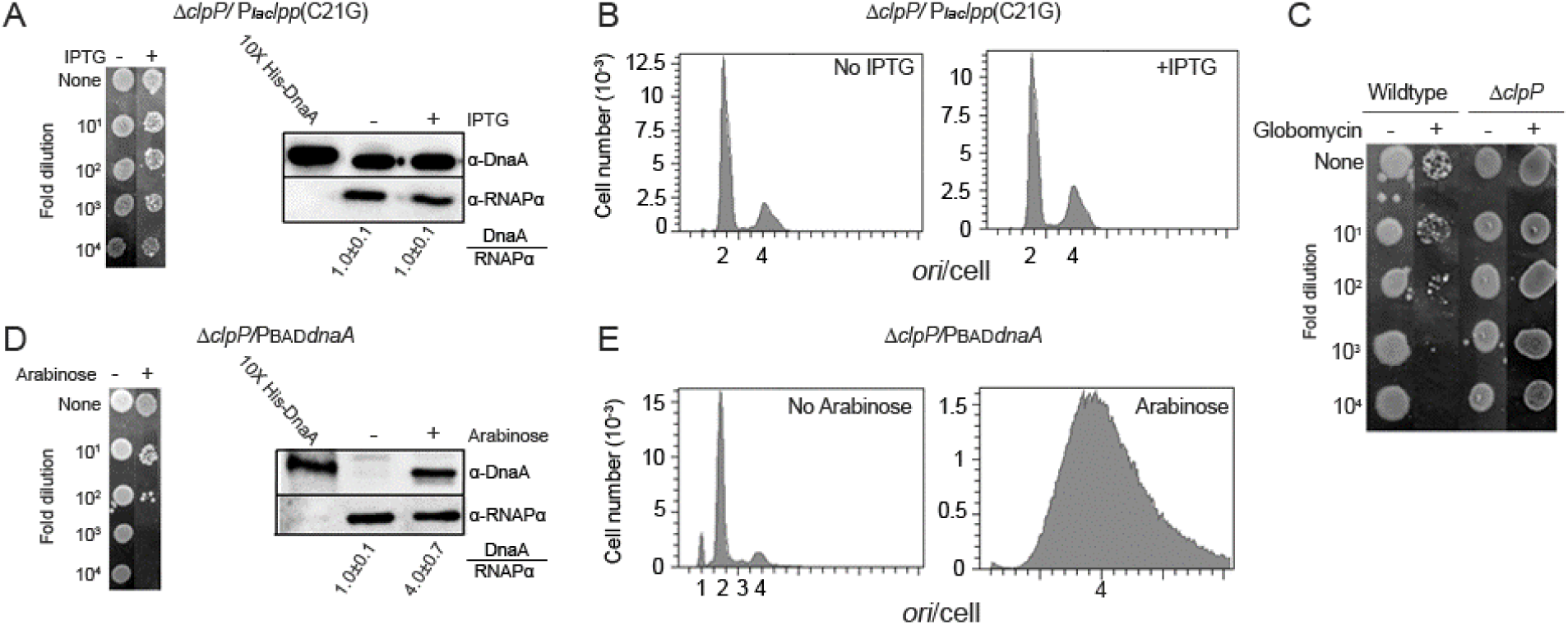
ClpP-dependence of DnaA depletion and replication blockage upon membrane-stress. (*A*) *E. coli* Δ*clpP* cells transformed with plasmid carrying the stressor gene P*_lac_lpp*(C21G) were grown without and with IPTG. Cell viability was assayed by spotting (*left panel*) and DnaA content of such cells by immunoblotting (*right panel*). Other details are same as in Fig. 4 A-B. (*B*) Replication run-off profile analyzed by flow cytometry of cells as in (*A*). (*C*) Growth of Δ*clpP* cells in the absence and presence of globomycin by spotting assay. (*D*) Growth (*left panel*) and DnaA content (*right panel*) of Δ*clpP* cells carrying the DnaA overexpressor plasmid. (*E*) Replication run-off profiles before and after induction of DnaA expression.

### Overexpression of *dnaA* overcomes the growth-arrest in Δ*crp* cells although not in wildtype cells

Although ClpP appears to be the protease that targets DnaA, how the membrane-stress leads to activation of the protease remains to be explored. Since we ruled out (p)ppGpp as the signaling molecule in the membrane-stress response (Fig. 4A), we considered the role of another common second messenger, cyclic (3′, 5′)-adenosine phosphate (cAMP), in downstream signaling of the membrane-stress that culminates in ClpP-mediated DnaA loss (Fig. 5).

We tested initially the role of CRP in the absence of inducing stress. The wildtype and Δ*crp* cells were transformed with plasmids carrying P_BAD_*dnaA* and only P_BAD_ (as a negative control). When DnaA synthesis was induced by adding arabinose, only Δ*crp* cells showed growth but not the wildtype cells (compare Fig. 6A and 6C, *lanes 1 vs. 2*). Immunoblotting indicated similar expression of DnaA in wildtype and Δ*crp* cells (Fig. 6A and C, *lanes 1 vs. 2*). Flow cytometric analysis of cells without the inducer showed roughly similar DNA content (with 2, 4, 6 or 8 chromosomes) in wildtype and Δ*crp* cells (Fig. 6B *vs.* 6D, no IPTG/no arabinose). When DnaA synthesis was induced, there was massive hyperinitiation in wildtype cells but not in Δ*crp* cells (Fig. 6B *vs.* 6D, no IPTG/+arabinose). These results indicate that the massive hyperinitiation could be responsible for inviability of wildtype cells as was seen earlier for Δ*lon* cells (Fig. 4F). The hyperinitiation was less in Δ*crp* cells and they were viable. Δ*crp* cells thus provided an opportunity to test whether overexpression of *dnaA* could overcome the growth-arrest upon induction of the membrane-stress.

**Fig. 6.**
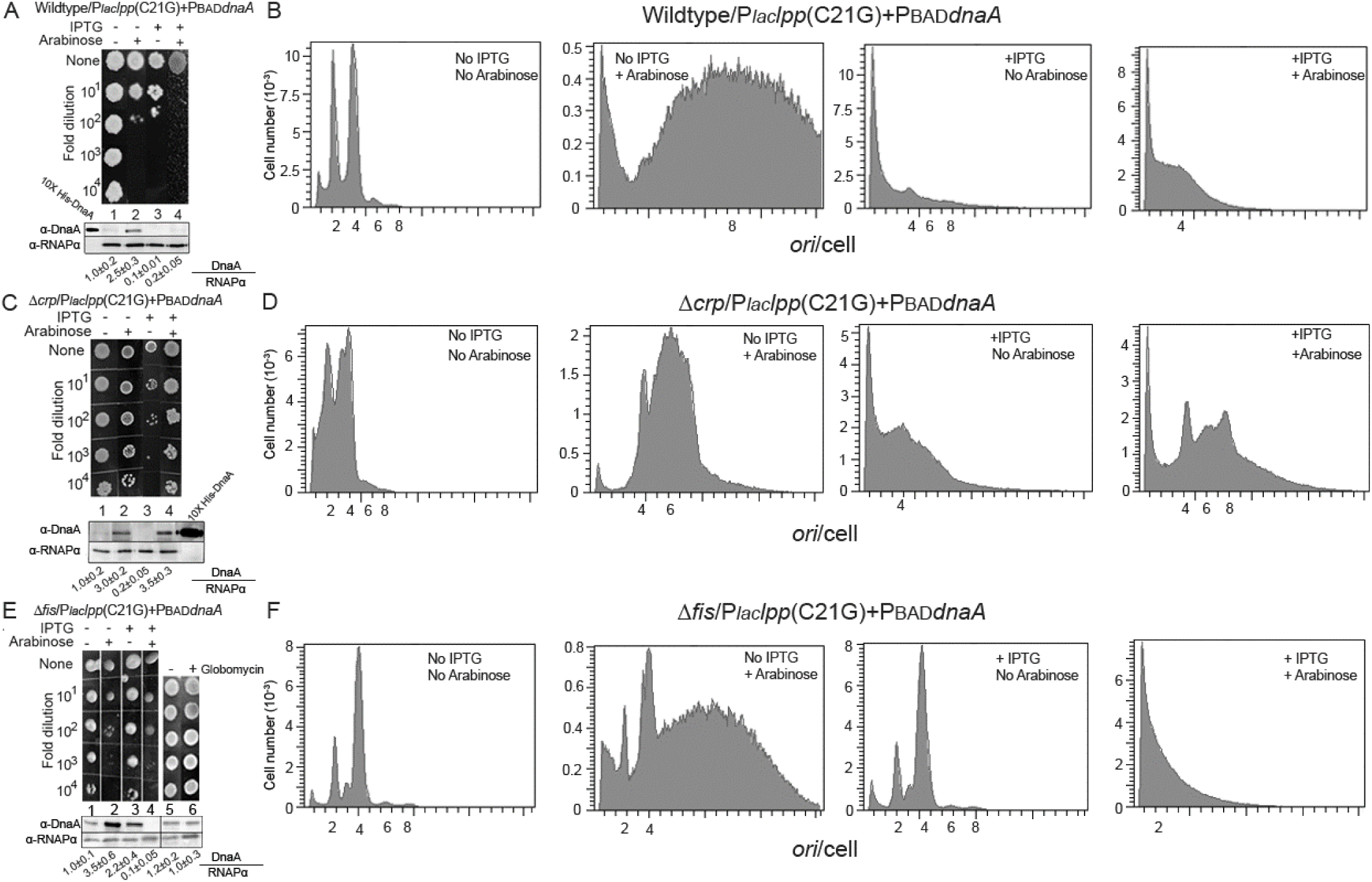
Overcoming of membrane-stress in Δ*crp* and Δ*fis* cells. *E. coli* wildtype, Δ*crp* and Δ*fis* cells were transformed with a single plasmid containing P_BAD_*dnaA* or in conjunction with another plasmid containing P*_lac_lpp*(C21G). *(A)* Effect of DnaA overexpression on growth of wildtype cells in the absence or presence of membrane-stress (i.e., ±IPTG) was tested by the spotting assay (*top panel*) and DnaA content of such cells by immunoblotting (*bottom panel*). *(B)* DNA content of the cells in (*A*) was tested by flow cytometry. *(C-D)* Same as *(A-B)* except that the cells were Δ*crp. (E-F)* Same as *(A-B)* except that the cells were Δ*fis.* In (*E*), growth of Δ*fis* cells was also tested for sensitivity to globomycin.

The wildtype and Δ*crp* cells containing the P_BAD_*dnaA* plasmid were further transformed with the plasmid containing P*_lac_lpp*(C21G), and the transformants were selected on plates without any inducer. Induction of *lpp*(C21G) (by IPTG inclusion) was inhibitory for both the wildtype and Δ*crp* cells (Fig. 6A *vs.* 6C, *lane 3*). In agreement, both the cells showed loss of DnaA by immunoblotting (Fig. 6A *vs.* 6C, *lane 3*), and replication block by flow cytometry (Fig. 6B *vs.* 6D, +IPTG). These results indicate that CRP by itself does not cause lethality from membrane-stress. However, when production of both DnaA and pLpp(C21G) was induced, only Δ*crp* and not the wildtype cells could overcome the growth-arrest (Fig. 6A *vs* 6C, *lane 4*). Immunoblotting of wildtype cells showed loss of DnaA (Fig. 6A, *lane 4*), which was also evident by reduced replication events in flow cytometric assays (Fig. 6B, +arabinose/+IPTG). In other words, the stress-induction is inducing a protease, which is depleting DnaA despite its overproduction. However, in Δ*crp* cells the DnaA level was not altered significantly upon the stress induction (Fig. 6B, *lanes 2 vs. 4*) and hyperinitiation was also modest (Fig. 6D, +arabinose/ +IPTG), explaining how cells could be viable. These results indicate that overcoming of the membrane-stress requires DnaA overproduction but toning it down to avoid hyperinitiation.

### DnaA is not lost and the growth is not arrested in Δ*fis* cells

We formerly found that cells absent the Fis (Factor for inversion stimulation) protein, which negatively regulates initiation from *oriC*, are not growth-arrested when they lack PG (Δ*pgsA*) or express the mutant *lpp*(C21G) gene (14, Fig. 6E, *lanes* 1 vs. 3). In line with these results, Δ*fis* cells when treated with globomycin also did not exhibit growth-arrest (Fig. 6E, *lanes 5 and 6*). Immunoblotting showed that DnaA is not lost in Δ*fis* cells upon induction of the membrane-stress with IPTG or upon treatment with globomycin (Fig. 6E, *lanes 1 vs. 3* and *5 vs. 6*). The flow cytometric data also agreed with the inference that upon induction of the membrane-stress, Δ*fis* cells are not blocked for replication initiation (Fig. 6F, *panel 1 vs 3*). Similarly, increasing DnaA content in Δ*fis* cells led to hyperinitiation-mediated growth inhibition as in wildtype cells (Fig. 6E, *lane 1* and Fig. 6F, 1^st^ *panel* vs. Fig. 6E *lane 2* and Fig. 6F, 2^nd^ *panel*). In addition, the membrane-stress in conjunction with DnaA overproduction led to growth-arrest due to DnaA loss and DNA degradation (Fig. 6E, *lane 4* and Fig. 6F, 4^th^ *panel*). These results demonstrate that Δ*fis* cells are resistant to membrane-stress like the Δ*crp* cells but without requiring *dnaA* overexpression.

## Discussion

Stress conditions such as nutrient starvation, fatty acid limitation, and antibiotic treatment accumulate <u>(p)ppGpp</u> molecules (47–49), which causes many regulatory changes, including activation of the Lon protease (50). Lon degrades, among others, DnaA and thus can block new rounds of replication (51, 52, 68, 69). Here, we find that induction of membrane-stress due to interrupted Lpp maturation activates the ClpP protease that degrades DnaA (Fig. 7). This prevents new rounds of replication initiation, which would suffice to explain cell-growth arrest upon the membrane-stress. This stress-response pathway appears novel as it is independent of several stress-response regulators such as Lon, DegP, and the alarmone (p)ppGpp. An independent report indicated that lethality due to interrupted trafficking of Lpp protein from IM to OM is mitigated in the absence of the two-component system CpxAR (75). Although no report has suggested any relationship between the activation of the CpxAR system and ClpP, our results suggest that the two could be causally related. The details of the pathway remain to be delineated, but Fis protein seems to be a requirement.

**Fig. 7.**
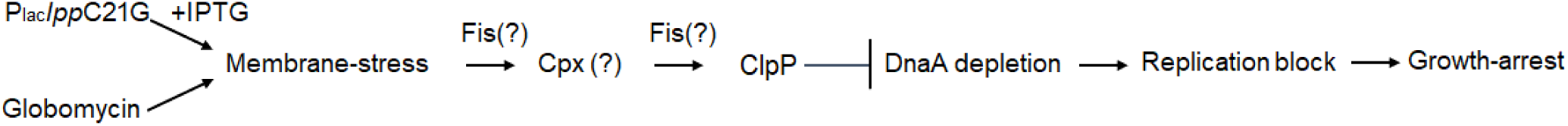
Requirement of Clp for growth-arrest upon membrane-stress. How ClpP is activated upon membrane-stress is not clear but a likely candidate is via the Cpx pathway. The requirement of Fis for the growth-arrest is more definite but at what stage of the stress-response pathway is unclear because the details of the pathway itself remains largely unknown, except that the Lon, DegP, (p)ppGpp and CRP are not required.

In the bacterium *E. coli* and *Salmonella enterica*, Fis is essential for regulating genes involved in stress-response pathways, particularly those related to antioxidant defense (76, 77). This function enables cells to effectively counteract any oxidative damage. Moreover, Fis plays a pivotal role in activating genes associated with nutrient uptake and metabolism, ensuring optimal resource utilization (78). We previously showed that the Lpp mutant pLpp(C21G), whose accumulation in the IM causes the membrane-stress in wildtype cells, is not accumulated in Δ*fis* cells, and these cells are not growth-arrested (14). In agreement, we find here that treatment with globomycin, which arrests growth of wildtype cells efficiently but not of Δ*fis* cells (Fig. 6E and F). This is expected since in the absence of the stressor pLpp(C21G) protein, inhibiting downstream players in the Lpp maturation pathway such as inhibiting LspA activity by globomycin should be inconsequential. Furthermore, we found no differences in the DnaA content and initiation frequency at the replication origin in Δ*fis* cells without and with membrane-stress. An independent Chip-Seq data indicated a specific reduction in ClpA levels in Δ*fis* cells compared to wildtype cells (79). This suggests that Fis might be controlling ClpA levels. We propose that in Δ*fis* cells, the reduced amount of ClpA might promote interaction of ClpP with its other AAA+ partner ClpX, the resultant increase in ClpXP formation would degrade the stressor pLpp(C21G), thereby enable normal cell-growth. The involvement of a global transcription factor like Fis in ClpP-mediated membrane stress, indicates that there are likely more genes within the stress-response pathway that are yet to be identified.

We also showed that membrane-stress upon defective lipoprotein biogenesis is evident equally in wildtype and Δ*crp* cells, meaning that CRP is unlikely to be a requirement for the growth-arrest. Since CRP function requires its cofactor cAMP, a well-known nucleotide second messenger, this suggests that the membrane-stress is not signaled through cAMP, analogous to the case with (p)ppGpp. We note that transcription of the *fis* operon sharply increases during the exponential growth phase, followed by a steep decrease as nutrients become scarce and the transcription ceases during the stationary phase (78). It is reported that both CRP and Fis are required to reduce the Fis levels in the cells (80). In other words, Fis level is expected to stay high in the absence of CRP (80). The elevated levels of Fis could keep increased ClpA levels and reduce the formation of ClpXP, leading to stability of pLpp(C21G) and growth-arrest. This scenario is fully consistent with our finding that Δ*crp* cells can be the growth-arrested (Fig. 6C and D). These cells, however, allowed us to confirm our central finding that the growth-arrest upon the membrane-stress is due to DnaA degradation since the arrest is overcome by overexpressing DnaA, which is lethal in wildtype cells but not in Δ*crp* cells. Apparently, the overexpression compensates for the DnaA-loss from the membrane-stress, allowing replication initiation and cell growth (Fig. 6C and D). In summary, although we have some understanding of the players in the stress-response pathway, much remains to be done to identify particularly the initial players of the pathway.

Our results suggest that globomycin although known as the direct inhibitor of the LspA enzyme, its bactericidal effect appears to be from DnaA proteolysis. A deeper understanding of the origin of membrane-stress and the stress-response that results in degrading DnaA is likely to provide further clues to prevent cell-growth, considering that DnaA is a universally conserved protein and vital to cell growth. The understanding can be of significant relevance to global health, considering that extended-spectrum β-lactamase-producing bacteria, such as *E. coli, Klebsiella pneumoniae, S. enterica*, and carbapenem-resistant *Enterobacteriaceae*, are prevalent in healthcare settings and becoming drug-resistant at an alarming rate (81, 82).

## Materials and Methods

We purchased restriction enzymes from New England Biolabs (NEB). The PCR primers used in this study were custom synthesized from Integrated DNA Technologies. We performed polymerase chain reactions using Q5 high-fidelity DNA polymerase (NEB). Chemical ingredients to make buffers and growth media were purchased from Sigma or VWR.

### Bacterial strains and plasmids

Bacterial strains and plasmids used in this study are mentioned in Supplementary Table S1.

#### Growth assay

Single transformants obtained by transforming wildtype (*pgsA^+^*) cells with plasmid DNA carrying *l*pp(WT), *lpp*(C21G) or *lpp*(C21G/ΔK) genes placed under the inducible P*_lac_* promoter were inoculated in M9 + Glu medium supplemented with ampicillin (100 μg/ml). The cultures grown overnight under non-inducing conditions, were diluted in M9+Glu medium to an OD_600_ of 0.005. The cultures were allowed to grow for an additional 45 minutes before dividing into two aliquots, with one containing isopropyl-β-D-galactopyranoside (50 µM IPTG) or globomycin (10 mg/ mL) antibiotic. The samples were collected at the indicated time intervals to monitor growth of cells carrying normal and mutant *lpp* genes.

#### Immunoblotting

At the last time point of the growth assay, the cells were pelleted by centrifugation at 16,000g for 10 min at 4 °C. Cell pellets were resuspended in 500 μl of PBS, and lysates were prepared by sonication for 5 minutes (in cycles of 10” off and 10” on). Following the determination of protein concentrations by Bradford assay, protein aliquots were stored at −80 °C until use. Lysates containing 5 µg of protein mixed with 1x SDS sample buffer were boiled and the proteins were resolved on 12% SDS-PAGE. Proteins transferred to PVDF membranes were immunoblotted with polyclonal α-DnaA antiserum, and treated with stabilized peroxidase-conjugated secondary antibody (Thermo Fisher Scientific). Blots were reprobed with monoclonal α-RNAP (alpha subunit), whose amounts served as loading controls. Immunoblots were visualized using AI600 Imager. The DnaA/RNAPa ratios are normalized w.r.t. the *lpp*(WT) value of 1.0. The experiments were performed as at least three biological replicates.

#### Spotting assay

The transformants were selected on M9-Agar supplemented with appropriate antibiotics but without IPTG. Single transformants were grown at 37 °C in M9+Glu media to an exponential phase (OD_600_ of 0.10). To perform the spotting assay, serial dilutions were prepared, and 2.5µL from each sample were placed on the M9-Agar+Glu media with or without Globomycin. Plates were allowed to grow at 37 °C for 16-24 hours.

#### Plating assay

*E. coli* WT *(pgsA^+^)* cells transformed with single or the two-plasmid DNA were selected on the M9-Agar+Glu containing plates supplemented with the appropriate antibiotics (ampicillin: 100 μg/ml, tetracycline: 12.5 μg/ml, kanamycin: 50 μg/ml). The viability of bacterial cells expressing DnaA(WT) and DnaA(L366K), alone or in conjunction with mutant Lpp(C21G), was tested. For this, single transformants containing either a single or two plasmid DNA were inoculated in M9+Glu media to an exponential phase (OD600 of 0.10). [Note that cells expressing pLpp(C21G) grow up to OD_600_ of 0.10 (Fig. 2A)]. Dilutions were prepared and plated to obtain 300-500 colonies on the M9-Agar plates containing the appropriate antibiotics, with or without IPTG, arabinose, or both.

#### Flow cytometry

Flow cytometry was performed as mentioned earlier (16). Briefly, overnight-grown cells were diluted to an optical density of ∼ 0.005 (@ 600 nm) in LB media supplemented with appropriate antibiotics. The cultures were divided into two equal volumes, to one of which IPTG (50 µM) was added to induce the expression of different *lpp* alleles. Cells were further grown to an OD_600_ of 0.15 and treated with rifampicin and cephalexin to inhibit new rounds of initiation and cell division, respectively. Cells grown for an additional 2.5 hours to allow completion of ongoing rounds of replication were subsequently analyzed for replication parameters at the NIH facility.

#### QRT-PCR

Cells grown in LB or M9 media were lysed at 4 ⁰C using disruptor beads and vortexing. Total RNA was extracted, and samples treated with RNase-free DNase to eliminate possible genomic DNA contamination. RNA concentration and integrity were assessed before performing cDNA synthesis. Each real-time PCR reaction will be limited to 40 cycles, and melt-curve analysis will be performed to ensure fidelity of the amplicon. Data were normalized to the ΔCq value of a commonly used reference gene, *rrsA* (16S rRNA gene), and the fold changes in the expression of the target genes were estimated.

## FUNDING AND ACKNOWLEDGEMENT

We want to acknowledge Dr. Ferenc Livak from the Flow Cytometry Core Facility, Center for Cancer Research, National Cancer Institute, National Institutes of Health, Bethesda, Maryland, USA. The Office of the Vice President of Academic and Faculty Affairs and the Office of the Dean of Research, Georgetown University Medical Center, provided financial support for this research.

